# Epstein-Barr Virus Induced Cytidine Metabolism Roles in Transformed B-cell Growth and Survival

**DOI:** 10.1101/2021.01.08.426018

**Authors:** Jin-Hua Liang, Chong Wang, Stephanie Pei Tung Yiu, Bo Zhao, Rui Guo, Benjamin E. Gewurz

## Abstract

Epstein-Barr virus (EBV) is associated with 200,000 cancers annually, including B-cell lymphomas in immunosuppressed hosts. Hypomorphic mutations of the *de novo* pyrimidine synthesis pathway enzyme cytidine 5’ triphosphate synthase 1 (CTPS1) suppress cell mediated immunity, resulting in fulminant EBV infection and EBV+ central nervous system (CNS) lymphomas. Since CTP is a critical precursor for DNA, RNA and phospholipid synthesis, this observation raises the question of whether the isozyme CTPS2 or cytidine salvage pathways help meet CTP demand in EBV-infected B-cells. Here, we found that EBV upregulated CTPS1 and CTPS2 with distinct kinetics in newly infected B-cells. While CRISPR CTPS1 knockout caused DNA damage and proliferation defects in lymphoblastoid cell lines (LCL), which express the EBV latency III program observed in CNS lymphomas, double CTPS1/2 knockout caused stronger phenotypes. EBNA2, MYC and non-canonical NF-□B positively regulated CTPS1 expression. CTPS1 depletion impaired EBV lytic DNA synthesis, suggesting that latent EBV may drive pathogenesis with CTPS1 deficiency. Cytidine rescued CTPS1/2 deficiency phenotypes in EBV-transformed LCL and Burkitt B-cells, highlighting CTPS1/2 as a potential therapeutic target for EBV-driven lymphoproliferative disorders. Collectively, our results suggest that CTPS1 and CTPS2 have partially redundant roles in EBV-transformed B-cells and provide insights into EBV pathogenesis with CTPS1 deficiency.

## Introduction

Epstein-Barr virus (EBV) causes infectious mononucleosis and persistently infects 95% of adults worldwide. For most individuals, lifelong EBV colonization of the memory B-cell compartment is asymptomatic. Nonetheless, EBV is associated with 200,000 human cancers per year, including Burkitt lymphoma, post-transplant lymphoproliferative diseases, Hodgkin lymphoma, gastric and nasopharyngeal carcinoma (1, 2). Endemic Burkitt lymphoma caused by EBV remains the most common childhood cancer in sub-Saharan Africa (3-5).

EBV is also associated with a growing number of rare congenital immunodeficiency syndromes, in which impairment of cell mediated immune responses disrupts the host/virus balance (6, 7). Inborn errors of immunity associated with severe EBV disease include *SH2D1A* and *XIAP* mutations, which cause the X-linked lymphoproliferative diseases 1 and 2 syndromes, respectively (8). Severe EBV infection is also associated with deficiency of the MAGT1 transporter, GATA2, the IL-2-inducible T-cell kinase, the RAS guanyl-releasing protein 1 RASGRP1, CD27, CD70 and 4-1BB (6, 7, 9-13). Chronic active EBV and EBV+ lymphomas occur with gain-of-function PI3 kinase catalytic subunit mutations that cause T-cell senescence (14).

Autosomal recessive mutations of the cytidine 5’ triphosphate synthetase 1 *CTPS1* gene cause a primary immunodeficiency syndrome characterized by impaired T-cell proliferation, low numbers of invariant and mucosal-associated T-cells and NK cells, and susceptibility to encapsulated bacterial and chronic viral infections, particularly by EBV and the related varicella zoster virus (15, 16). Most CTPS1 deficient patients experienced EBV disease, including severe infectious mononucleosis, chronic EBV viremia and EBV-associated primary CNS lymphomas (16). Extra-hematopoietic manifestations of CTPS1 deficiency have not been reported, and hematopoietic stem cell transplantation is curative (17).

The nucleotide cytidine is a key precursor for DNA, RNA and phospholipid biosynthesis as well as protein sialyation. Cytidine triphosphate (CTP) is maintained at the lowest concentration of cellular nucleotide pools and its synthesis is tightly regulated (18, 19). CTP is produced through *de novo* or salvage pathways. The isozymes CTPS1 and CTPS2 catalyze the ATP-dependent transfer of nitrogen form glutamine to uridine triphosphate (UTP), thereby producing CTP and glutamate. CTPS1 and 2 share 74% amino acid sequence identity (20, 21). CTP can also be synthesized by salvage pathways, where uridine or cytidine are imported and phosphorylated by the enzymes uridine-cytidine kinase 1 or 2 (UCK1 or UCK2). Cytidine monophosphate is subsequently converted to CTP by the kinases CMPK and NDPK (**Figure 1a**). CTP synthases are the rate-limiting enzymes in *de novo* CTP synthesis (22), and CTPS1 is rapidly and highly upregulated in primary human T-cells upon T-cell receptor stimulation (23).

**Figure 1.**
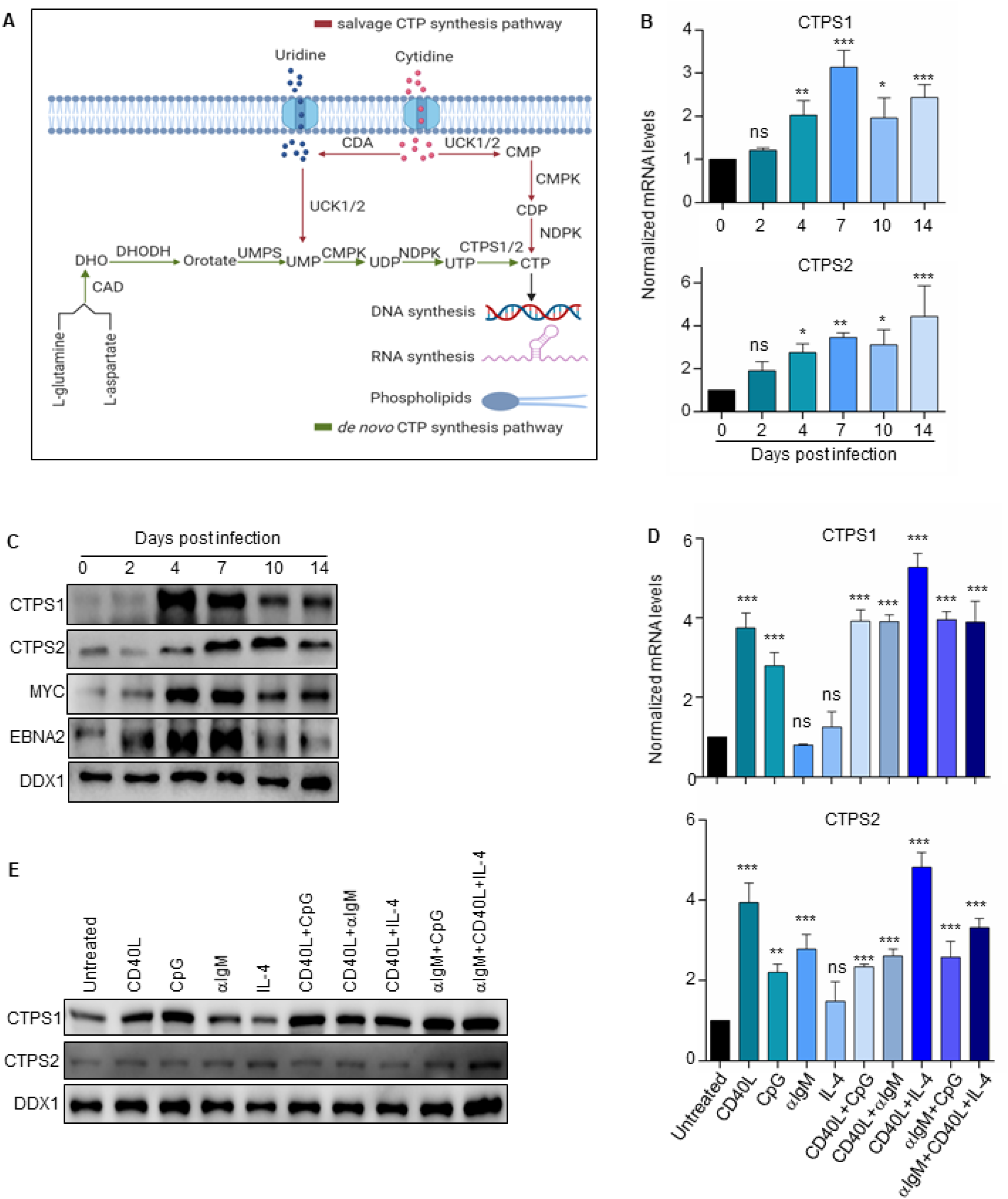
Upregulation of *de novo* CTP synthesis enzymes CTPS1 and CTPS2 in primary human B-cells stimulated by EBV or B-cell agonists. **(A)** Schematic diagram of *de novo* and salvage CTP synthesis pathways. The *de novo* pathway synthesizes CTP from L-glutamine and L-aspartate or uridine substrates, whereas the salvage pathway uses cytidine for CTP synthesis. **(B)** Quantitative PCR (qPCR) analysis of CTPS1 and CTPS2 mRNA levels in primary human CD19+ peripheral blood B-cells infected by EBV B95.8 strain for the indicated days. **(C)** Immunoblot analysis of CTPS1, CTPS2, MYC, EBNA2 and load control DDX1 levels in whole cell lysate (WCL) from primary human CD19+ peripheral blood B-cells infected by the EBV B95.8 strain for the indicated days. **(D)** qPCR analysis CTPS1 and CTPS2 mRNA levels in primary human CD19+ B-cells stimulated for 24 hours by Mega-CD40 ligand (CD40L, 50ng/mL), the toll-receptor 9 agonist CpG(1μM), □IgM crosslinking (1 μg/mL), interleukin 4 (IL-4, 20ng/mL) or combinations thereof, as indicated. (**E**) Immunoblot analysis of CTPS1, CTPS2 and DDX1 abundances in WCL of primary human CD19+ B-cells stimulated for 24 hours by CD40 ligand Meda-CD40L (50ng/mL), CpG (1μM), □IgM (1μg/mL), IL-4 (20ng/mL), or combinations thereof, as indicated. For d and d, mean and SEM values from n=3 replicates are shown. *, *p*<0.05; **, *p*<0.01; ***, *p*<0.001; ****, *p*<0.0001; ns, non-significant using one way Anova with multiple comparisons. For c and d, representative blots from n=3 replicates are shown.

All patients described thus far with CTPS1 deficiency harbor a homozygous missense G-> C mutation at base 1692 of the *CTPS1* gene (rs145092287) that alters an intronic splice acceptor site. While the mutation does not alter CTPS1 catalytic activity, it causes 80-90% loss of CTPS1 protein abundance (16). The high prevalence of B-cell driven EBV-associated diseases in people with CTPS1 deficiency raises the question of how EBV-driven lymphomas arise at high frequency, despite pronounced defects in lymphocyte metabolism. Patients with CTPS1 deficiency have normal total T and B-cell numbers, though reduced frequency of memory B-cells, the site of long-term EBV persistence (16). Whether CTPS2 may have more important roles in EBV-infected B-cells than in T and NK cells, and how CTPS1 deficiency alters the latent versus lytic stages of the EBV lifecycle remain unknown.

B-cells from individuals with CTPS1 deficiency exhibit defects in proliferation (16, 23). Furthermore, the *de novo* pyrimidine synthesis enzyme DHODH inhibitor leflunomide impairs growth of EBV-driven lymphomas in murine models (24). Nonetheless, continuously growing EBV-transformed lymphoblastoid cell lines (LCLs) can be established from B-cells of patients with CTPS1 deficiency (16, 23). LCLs express the EBV latency III program also observed in primary CNS lymphomas, comprised by six Epstein-Barr nuclear antigens (EBNA) and two latent membrane proteins (LMP) that mimic signaling by the B-cell receptor and CD40 (25, 26).

To gain insights into EBV pathogenesis in the setting of CTPS1 deficiency, we characterized roles of CTPS1, CTPS2 and cytidine salvage pathways in support of EBV-transformed B-cell growth and lytic replication.

## Results

### EBV upregulates CTPS1 in newly infected B cells

To gain insights into EBV effects on CTPS1 and CTPS2 expression, we profiled primary human B-cells at rest and at five timepoints post infection. Whereas low levels of CTPS1 or CTPS2 were detected in resting peripheral blood CD19+ B-cells, EBV markedly upregulated both CTPS1 and CTPS2 mRNAs. CTPS1 mRNA reached maximum levels between days 4-7, the period where transforming cells undergo Burkitt-like hyper-proliferation (27, 28). By contrast, CTPS2 was progressively upregulated as newly infected cells converted to lymphoblastoid physiology between days 7-14 post-infection (**Figure 1b**). Similar effects were evident on the protein level (**Figure 1c**), where CTPS1 exhibited comparable expression patterns to Epstein-Barr nuclear antigen 2 (EBNA2) and MYC, each of which are important metabolism regulators. Using our recently published RNAseq and proteomic maps of EBV-driven B-cell growth transformation (28, 29), EBV was also found to upregulate nearly all components of the pyrimidine *de novo* and salvage pathways at similar timepoints (**Figure S1**).

CTPS1 and 2 are upregulated in primary B cells activated by combinatorial mitogenic stimuli, including by CD40-ligand (CD40L) plus interleukin-4 (IL-4), but responses to individual stimuli have not been studied (23). To cross-compare immune receptor and EBV effects on CTPS1 and CTPS2 expression, CD19+ primary B-cells were stimulated for 24 hours with CD40L, the Toll-like receptor 9 agonist CpG, □IgM, the cytokine IL-4, or combinations thereof (**Figure 1d-e**). Interestingly, CD40L or CpG alone were sufficient to upregulate CTPS1 mRNA and protein to near maximal levels, whereas CTPS2 was significantly induced on the mRNA but not protein level in response to most B-cell agonists (**Figure 1d-e**). These results highlight dynamic CTPS1 and CTPS2 regulation that varies over phases of EBV-mediated B-cell transformation and in response to particular receptor agonists.

### CTPS1 is important for EBV-infected B-cell outgrowth

Since LCLs from CTPS1 deficient patients retain 10-15% CTPS1 expression, this residual CTPS1 activity may be necessary for EBV-transformed B-cell growth or survival. Alternatively, CTPS2 redundancy may support these functions. To investigate these possibilities, we first re-analyzed data from our recent human genome-wide CRISPR/Cas9 growth and survival screen (30). Strong selection against all four single guide RNAs (sgRNAs) targeting the *CTPS1* gene was evident at screen day 21 of the LCL GM12878 and the EBV+ Burkitt cell line P3HR-1. By contrast, sgRNAs against *CTPS2* or the genes encoding pyrimidine salvage pathway enzymes UCK1 or UCK2 were not strongly selected against, suggesting that functional knockout of their targets were tolerated by these EBV-transformed B-cell lines (**Figure 2a**). Next, CRISPR mutagenesis was performed with *CTPS1* targeting sgRNAs to characterize acute CTPS1 loss-of-function effects on Burkitt and LCL proliferation (**Figure 2b**). Live cell numbers were measured at five timepoints following expression of control versus independent *CTPS1* targeting sgRNAs and puromycin selection. CTPS1 depletion significantly impaired growth of the EBV+ Burkitt cell line Daudi, which has latency I expression, and of the LCL GM12878 (**Figure 2b**). CTPS1 editing also diminished proliferation of GM11830 LCLs and Mutu I Burkitt cells (**Figure S2a-b**). A low level of proliferation was nonetheless observed in CTPS1 depleted cells, suggesting alternative routes of CTP synthesis.

**Figure 2.**
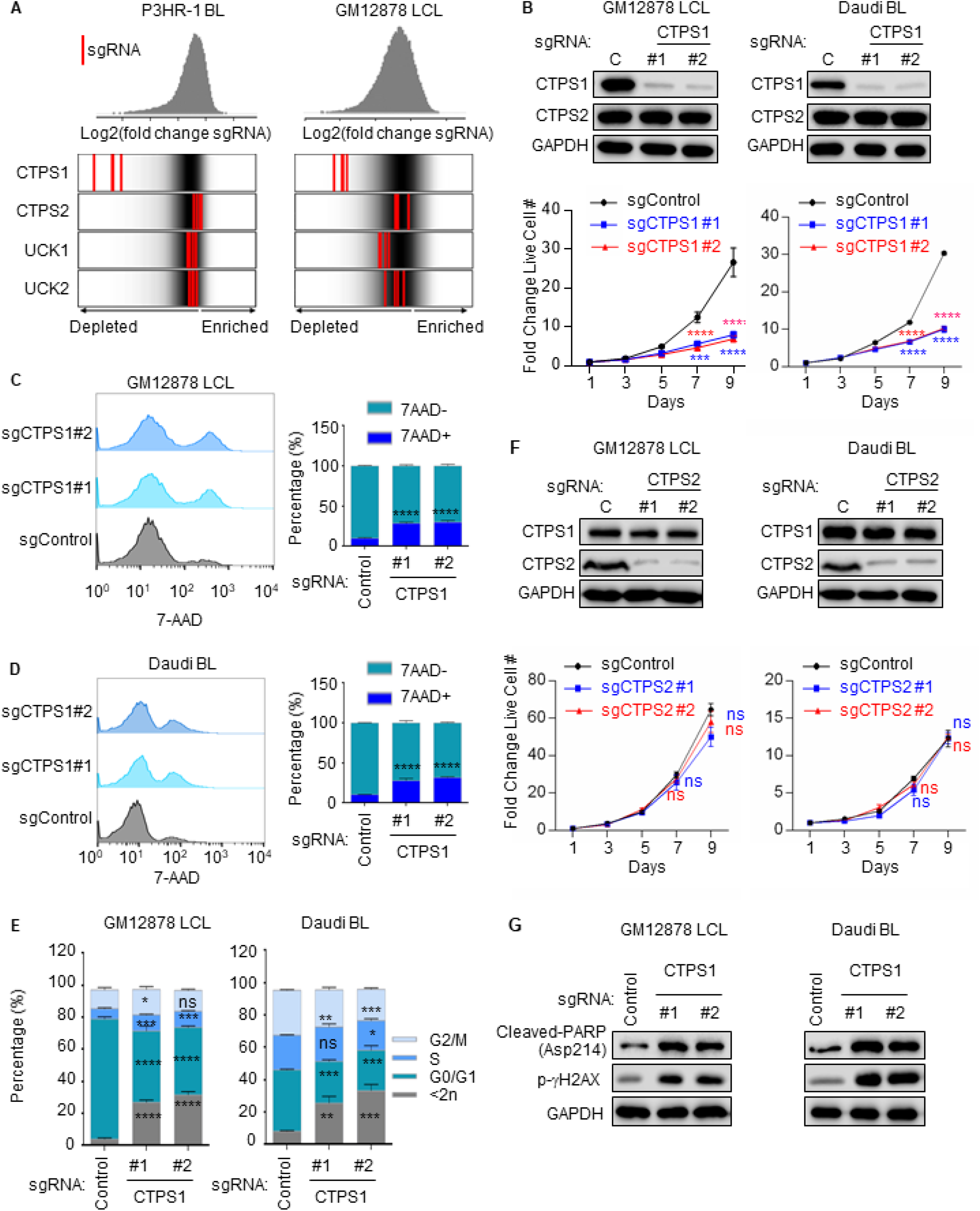
CTPS1 roles in EBV transformed B-cell growth and survival. **(A)** Distribution of Avana human genome-wide CRISPR screen sgRNA Log2 fold-change values at Day 21 versus input in Cas9+ P3HR-1 Burkitt lymphoma (BL) left or GM12878 LCL (right). Values for CTPS1, CTPS2, UCK1 or UCK2 targeting sgRNAs (red lines) are overlaid on gray gradients depicting all Avana sgRNA library values (30). Average Day 21 vs input values from four screen biological replicates are shown. (**B**) Immunoblot analysis (top) and growth curve analysis (bottom) of Cas9+ GM12878 LCLs or Daudi Burkitt lymphoma (BL) expressing non-targeting control (C) or independent sgRNAs against *CTPS1*. WCL were obtained at day 7 post sgRNA expression. Growth curves were begun at 96 hours post-sgRNA expression and 48 hours post-puromycin selection. **(C)** Flow cytometry (FACS) analysis of vital dye 7-AAD uptake in GM12878 LCL at Day 7 post-expression of control or sgRNAs targeting *CTPS1*. Shown at right are median + SEM values from n=3 replicates. (**D**) FACS analysis of 7-AAD uptake in Daudi BL at Day 7 post-expression of control or *CTPS1* targeting sgRNAs. Shown at right are median + SEM values from n=3 replicates. (**E**) Propidium iodide cell cycle analysis of GM12878 LCL (left) or Daudi BL (right) expressing control or CTPS1 sgRNAs at Day 7. (**F**) Immunoblot (top) and growth curve analysis (bottom) of GM12878 or Daudi expressing control or independent sgRNAs against CTPS2. WCL were obtained at day 5 post sgRNA expression. Growth curves were begun at 96 hours post-sgRNA expression and 48 hours post-puromycin selection. (**G**) Immunoblot analysis of WCL from GM12878 or Daudi, 7 days post expression of control or CTPS1 sgRNAs. For B, C, D, E and F, mean and SEM values from n=3 replicates are shown. *, *p*<0.05; **, *p*<0.01; ***, *p*<0.001; ****, *p*<0.0001; ns, non-significant using one way Anova with multiple comparisons. For B, F and G, representative blots or n=3 replicates are shown.

To examine effects of CTPS1 depletion on B-cell survival, 7-AAD vital dye exclusion assays were performed on LCLs and Burkitt cells 5 days post sgRNA expression and puromycin selection, the timepoint at which growth curves of control and CTPS1 depleted cells diverged. Loss of CTPS1 significantly increased 7-AAD uptake in GM12878 and Daudi cells (**Figure 2c-d**) and in a second LCL and Burkitt pair (**Figure S2c-d**), indicating induction of cell death. Cell cycle analysis demonstrated that CTPS1 editing also impaired cell growth, with arrest most pronounced at the G1/S stage (**Figure 2e and Figure S2e-f**). Depletion of CTPS1 induced phosphorylation of H2AX at Ser 139 (□-H2AX), indicative of DNA damage, perhaps results from imbalance in CTP nucleotide pools. PARP cleavage was also evident, indicating apoptosis induction (**Figure 2g, Figure S2g**).

### Compensatory CTPS2 role in EBV-transformed B-cell outgrowth

EBV+ CNS lymphomas are often observed in patients with CTPS1 deficiency, which are estimated to have 10-20% residual CTPS1 activity (16). Yet, the CRISPR Cancer Dependency Map (DepMap) (31) found that targeting of CTPS2 only mildly affects proliferation of a broad range of cancer cells, including B-cell lymphomas (**Figure S3a**). These observations raise the question of the extent to which CTPS2 can be recruited to support *de novo* salvage CTP synthesis in EBV-transformed B-cells in the context of hypomorphic CTPS1 activity.

To address this question, we next tested the effects of CTPS2 editing in LCLs and Burkitt cells, alone or in the context of CTPS1 depletion. In contrast to CTPS1 depletion, CTPS2 targeting did not significantly impair LCL or Burkitt B-cell growth or survival (**Figure 3a-d, Figure S3b-d**). However, concurrent CTPS1/2 targeting nearly completely abrogated LCL outgrowth, to a greater extent than depletion of CTPS1 alone (**Figure 3a-d, Figure S3e-f**). Cytidine supplementation rescued outgrowth of CTPS1 or CTPS1/2 doubly deficient GM12878 LCLs or Daudi Burkitt cells (**Figure 3e-h**), suggesting that CRISPR effects were on target and demonstrating that the cytidine salvage pathway was capable of restoring sufficient CTP pools. Interestingly, withdrawal of exogenous cytidine at Day 7 of the proliferation assay rapidly blocked outgrowth of CTPS1 deficient and CTPS1/2 doubly-deficient GM12878 and Daudi cells (**Figure 3i-l**), further underscoring their acquired dependence on salvage CTP metabolism. CTPS1/2 depletion caused DNA damage, apoptosis induction and cell death in GM12878 and Daudi, as judged by immunoblot for cleaved-PARP, phospho □H2AX signal (**Figure 3m**). 7-AAD uptake was higher following dual CTSP1/2 targeting (**Figure 3n**).

**Figure 3.**
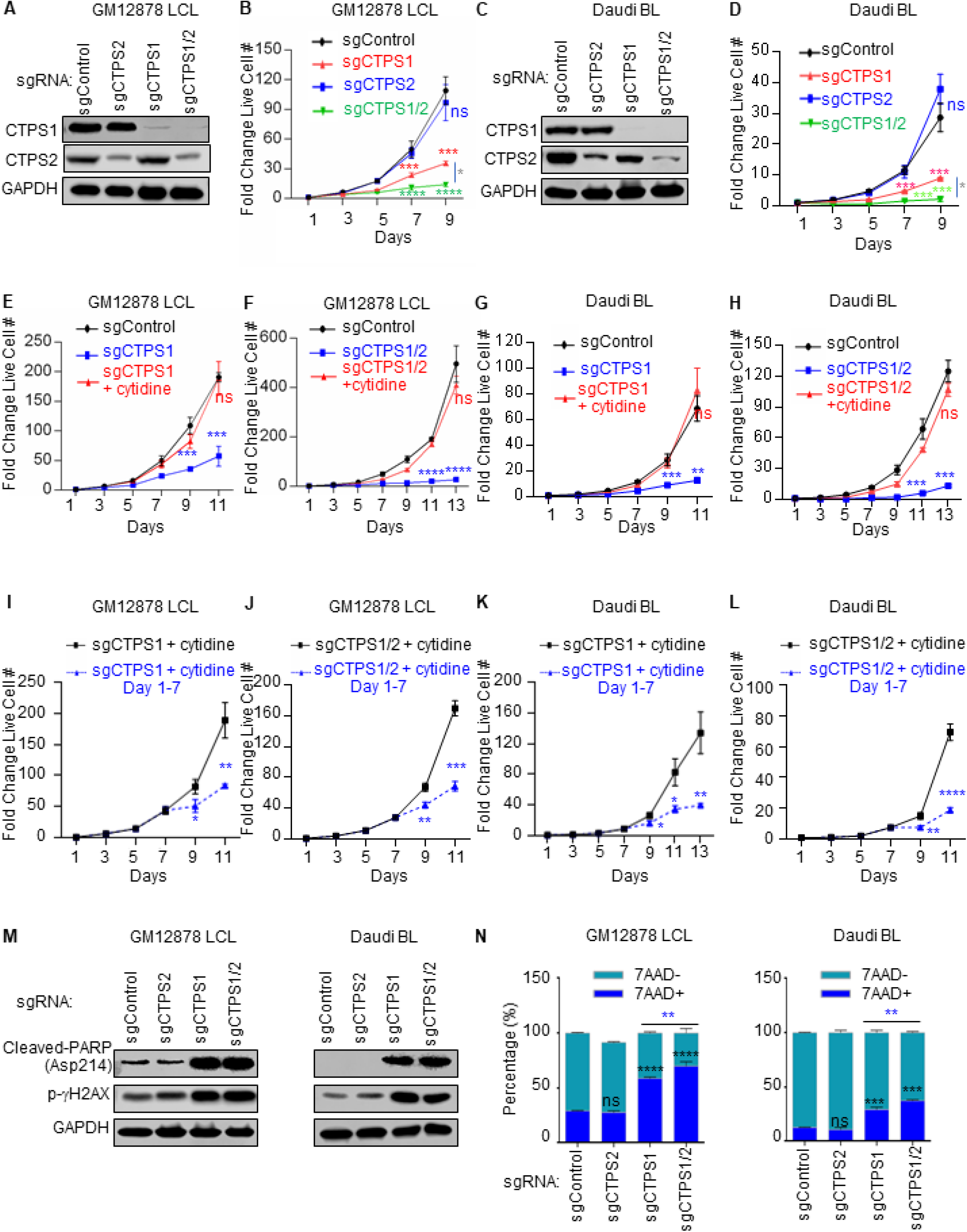
Partially redundant CTPS1/2 roles in EBV-transformed B cell growth and survival. **(A)** Immunoblot of WCL from GM12878 LCL on Day 7 post expression of the indicated control, CTPS1 and/or CTPS2 sgRNAs. **(B)** Growth curve analysis of GM12878 LCL with control, CTPS1 and/or CTPS2 sgRNAs, as indicated. **(C)** Immunoblot of WCL from Daudi BL Day 7 post expression of the indicated control, CTPS1 and/or CTPS2 sgRNAs. **(D)** Growth curve analysis of Daudi BL with control, CTPS1, and/or CTPS2 sgRNAs, as indicated. **(E-H)** Growth curve analysis of GM12878 LCL (E-F) or Daudi BL (G-H) that express control, CTPS1 or CTPS1 and CTPS2 sgRNAa as indicated, with 200 □M cytidine supplementation. **(I-L)** Growth curve analysis of GM12878 (I), GM11830 (J), Daudi (K) or Mutu I (L) with CTPS1 sgRNA expression, cultured in the presence of 200 □M cytidine from days 1-7 or throughout. **(M)** Immunblot analysis of WCL from GM12878 (left) or Daudi (right) at Day 7 post expression of the indicated sgRNAs. (**N**) Mean + SEM values of FACS 7-AAD values from GM12878 (left) or Daudi (right) expressing the indicated sgRNAs for 7 days from n=3 replicates. Blots in A, C, and M are representative of n=3 replicates. Growth curves were begun at 96 hours post-sgRNA expression and 48 hours post-puromycin selection. Growth curves show mean + SEM fold change live cell numbers from n = 3 replicates. *, *p*<0.05; **, *p*<0.01; ***, *p*<0.001; ****, *p*<0.0001; ns, non-significant using one way Anova with multiple comparisons.

### CTPS1 Transcription upregulation in EBV-transformed B-cells

The T-cell receptor and ERK pathways are important for rapid CTPS1 expression in activated T-cells. As pathways that control B-cell CTPS1 induction remain uncharacterized, we next investigated factors important for *CTPS1* expression in EBV-transformed B-cells. First, we used our and ENCODE published LCL ChIP-seq datasets to characterize occupancy of EBV nuclear antigens and EBV LMP1-activated NF-□B tranxcription factors (32-34). This highlighted *CTPS1* promoter region occupancy by EBV transcription factor EBNA-LP, by multiple NF-□B transcription factor subnits, by c-MYC and MAX (**Figure 4a**). Interestingly, EBNAs 2, LP, 3A, 3C and NF-□B subnits ReIB and p52 co-occupied an intergenic region ∼12 kilobases downstream from the *CTPS1* promoter. RNA pol II GM12878 Chromatin Interaction Analysis by Paired-End Tag Sequencing (ChIA-PET) analysis identifies a putative long-range interaction between Pol II bound to this *CTPS1* intragenic region and the *CTPS1* promoter (35).

**Figure 4.**
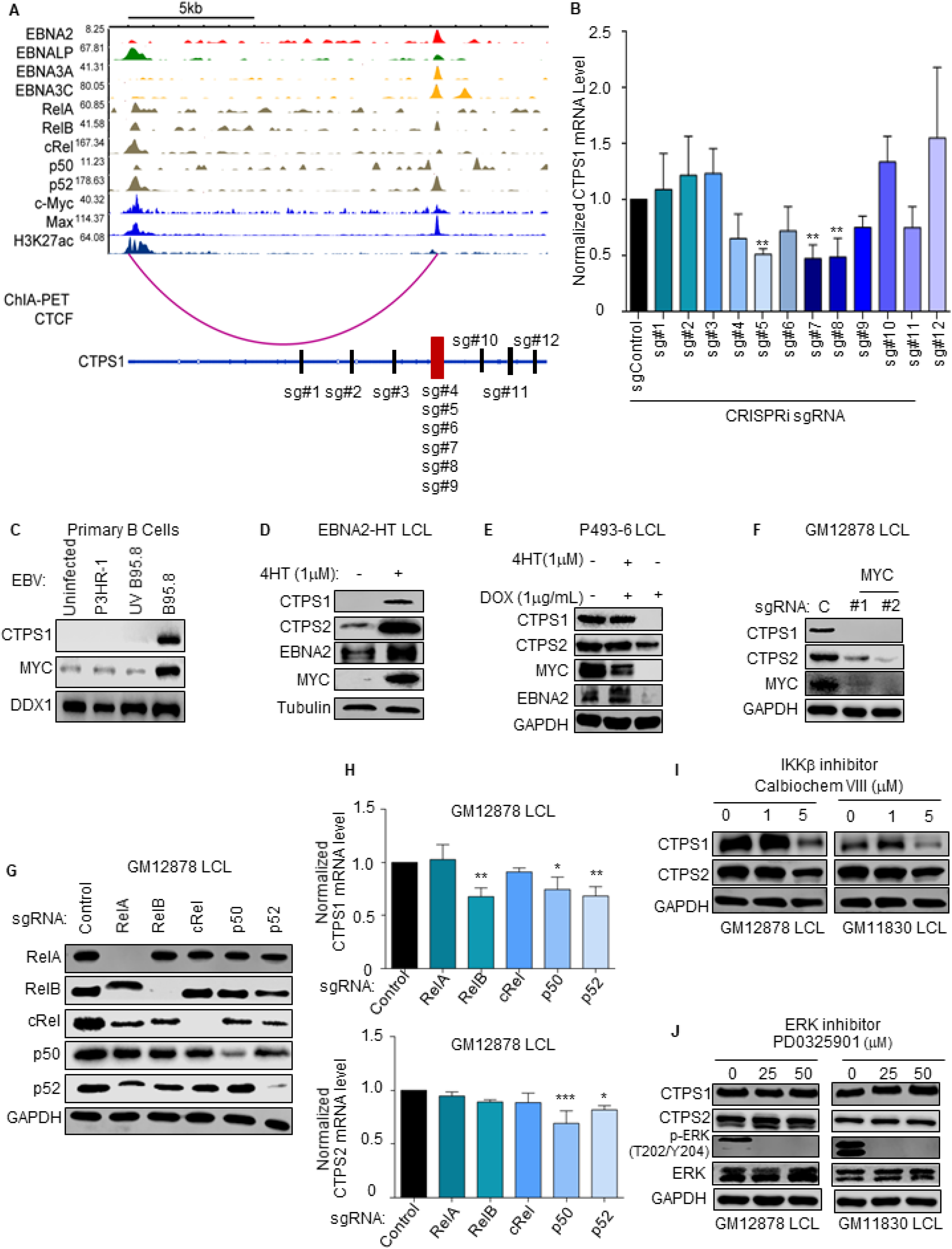
CTPS1 Transcription Regulation in EBV-infected B-cells. **(A)** Chromatin immunoprecipitation (ChIP)-sequencing (ChIP-seq) tracks for the indicated transcription factors or histone 3 lysine 27 acetyl (H3K27Ac) at the LCL *CTPS1* locus. Y-axis ranges are indicated for each track. Also shown below is LCL RNA Pol II ChIA-PET signal indicating a long-range DNA interaction between the *CTPS1* promoter region and an intronic site. Schematic diagram of the *CTPS1* locus shown below, with exons indicated by rectangles. CRISPR-i sgRNA targeting sites are shown. **(B)** Normalized CTPS1 mRNA levels in dCAS9-KRAB+ GM12878 expressing the indicated sgRNAs for CRISPR-i analysis for 5 days. CTPS1 mRNA level in cells with non-targeting control sgRNA was set to 1. **(C)** Immunoblot of WCL from primary human B-cells that were either mock-infected or infected with equal amounts of the non-transforming P3HR-1, UV-irradiated B95-8 or B95-8 EBV strain for four days. **(D)** Immunoblot analysis of samples prepared from 2-2-3 EBNA2-HT LCLs grown in the absence (EBNA2 non-permissive) or presence (EBNA2 permissive) of 4HT (1 μM) for 48 hours. **(E)** Immunoblot analysis of WCL from conditional P493-6 LCLs treated with 1 ug/ml doxycycline (DOX) to suppress exogenous *MYC* allele expression and/or with 1 M 4HT to induce EBNA2 activity for 48 hours, as indicated. **(F)** Immunoblot analysis of WCL from GM12878 LCL expressing control or independent *MYC*-targeting sgRNAs for 5 days, as indicated. **(G)** Immunoblot analysis of WCL from GM12878 LCL expressing the indicated sgRNA for 6 Days. (**H**) qPCR analysis of CTPS1 and CTPS2 mRNA levels in GM12878 LCL expressing the indicated sgRNAs for 7 days. Levels in cells with sgControl were set to 1. (**I**) Immunoblot of WCL from GM12878 or GM11830 LCLs treated with the indicated concentrations of the IKK antagonist Calbiochem VIII for 48 hours. 1 □M selectively blocks the canonical pathway IKK□ kinase. (**J**) Immunoblot of WCL from GM12878 or GM11830 LCLs treated with the ERK inhibitor PD0325901 at the indicated concentrations for 48 hours. Blots in C, D, E, F, G, I and J are representative of n = 3 experiments. SEM + Mean values are shown in B and H from n=3 replicates. *, *p* < 0.05; **, *p* < 0.01; ***, *p*<0.001 using one way Anova with multiple comparisons.

To study the functional significance of the EBNA and NF-□B-bound *CTPS1* intragenic region in *CTPS1* transcription regulation, we used CRISPR-interference (CRISPR-i) (36). GM12878 LCLs with endonuclease dead Cas9 fused to a KRAB transcription repressor domain were transduced with lentiviruses that express control sgRNA versus one of the twelve sgRNAs indicated in **Figure 4a**. Following puromycin selection, CRISPR-i effects on steady state *CTPS1* mRNA abundance were quantified by real time PCR. Four of the six sgRNAs targeting the putative enhancer significantly reduced CTPS1 mRNA levels. By contrast, none of the six sgRNAs targeting upstream or downstream control regions diminished CTPS1 mRNA (**Figure 4a-b**). These results suggest that the EBV latency III growth program induces *CTPS1* induction through effects at the *CTPS1* promoter and at a downstream interacting region.

EBNA2 and EBNA-LP are the first EBV proteins expressed in newly infected cells. To gain insights into potential EBNA2 roles in EBV-driven *CTPS1* induction, we infected human peripheral blood CD19+ B-cells, isolated by negative selection, with transforming B95.8 EBV, with ultraviolet (UV) inactivated B95.8, or with non-transforming P3HR-1 EBV that lacks EBNA2 and part of the EBNA-LP open reading frames. Only infection by B95.8 induced MYC, a well characterized regulator of metabolism, as well as CTPS1 by 48 hours of infection (**Figure 4c**). These data suggest that EBNA2 and/or EBNA-LP, rather than an innate immune response to the incoming viral particle, are required for CTPS1 induction in newly infected B-cells. We next used 2-2-3 LCLs with conditional EBNA2 expression, in which EBNA2 is fused to the ligand binding domain of a modified estrogen receptor that binds to 4-hydroxy-tamoxifen (4HT), to study EBNA2 roles in LCL CTPS1 expression. When grown in non-permissive conditions in the absence of 4HT for 48 hours, CTPS1 expression was rapidly lost, as was expression of EBNA2 target gene MYC (**Figure 4d**). Thus, EBNA2, perhaps together with its host target genes, are important for EBV-driven CTPS1 induction.

To characterize putative CTPS1 induction roles of the key host EBNA2 target MYC, we used p493-6 LCLs, which carry conditional 4HT-controlled EBNA2 and tetracycline-repressed *MYC* alleles (37). EBNA2 inactivation by 4HT withdrawal again caused loss of CTPS1 and MYC expression in cells cultured with doxycycline to suppress exogenous *MYC* (**Figure 4e**). Interestingly, induction of exogenous MYC expression by doxycycline withdrawal restored CTPS1 expression in the absence of EBNA2 activity (**Figure 4e**). Similarly, MYC knockout in GM12878 blocked CTPS1 expression at a timepoint prior to induction of cell death (**Figure 4f**), further supporting a key role for MYC in CTPS1 regulation and providing a potential route for CTPS1 induction in Burkitt cells.

The EBV-encoded CD40 mimic LMP1 activates the NF-□B canonical and non-canonical pathways to trigger nuclear translocation of the five NF-□B transcription factor subunits. Since GM12878 ChIP-seq identified *CTPS1* locus occupancy by all NF-□B subunits, we used CRISPR to individually test their roles in LCL CTPS1 expression (**Figure 4g**). Depletion of RelB, p50 and p52 each significantly reduced CTPS1 steady-state mRNA levels (**Figure 4h**), implicating the non-canonical NF-□B pathway. In support,a □B kinase (IKK)-□-selective antagonist did not visibly reduce CTPS1 protein levels at a dose that also blocks the kinaes IKK□, where the canonical pathway is selectively blocked, but impaired CTPS1 expression at a higher dose which controls the non-canonical pathway (**Figure 4i**). By contrast, ERK inhibition by an antagonist that blunts T-cell CTPS1 expression (23) failed to reduce CTPS1 expression, in spite of LMP1’s ability to activate ERK (**Figure 4j**). Thus, distinct mechanisms that control CTPS1 expression in EBV+ B-cells versus T-cells.

### EBV+ B-cells use *de novo* and salvage UTP metabolism

Uracil monophosphate (UMP), a key precursor in *de novo* CTP synthesis, can be synthesized from L-glutamine and L-aspartate by the sequential enzymatic activities of CAD, DHODH and UMPS. UMP can also be produced by UCK1 or UCK2 phosphorylation of imported uridine (**Figure 1a**). Interestingly, UCK was originally identified as a B-cell factor that associates with EBNA3A, suggesting possible roles in EBV-driven B-cell proliferation (38). To gain insights into potential DHODH versus UCK1/2 roles in newly infected primary B-cells, immunoblot analysis was performed on extracts from cells prior to and at five timepoints post-EBV infection. UCK2 was robustly induced by day 4, whereas DHODH expression was suppressed by day 10, despite each being induced on the mRNA level (**Figure 5a, Figure S4a-b**). By comparison, UCK1, 2 and DHODH mRNAs were upregulated by multiple primary B-cell stimuli, while again UCK2 but not DHODH was upregulated on the protein level (**Figure 5b, Figure S4c)**.

**Figure 5.**
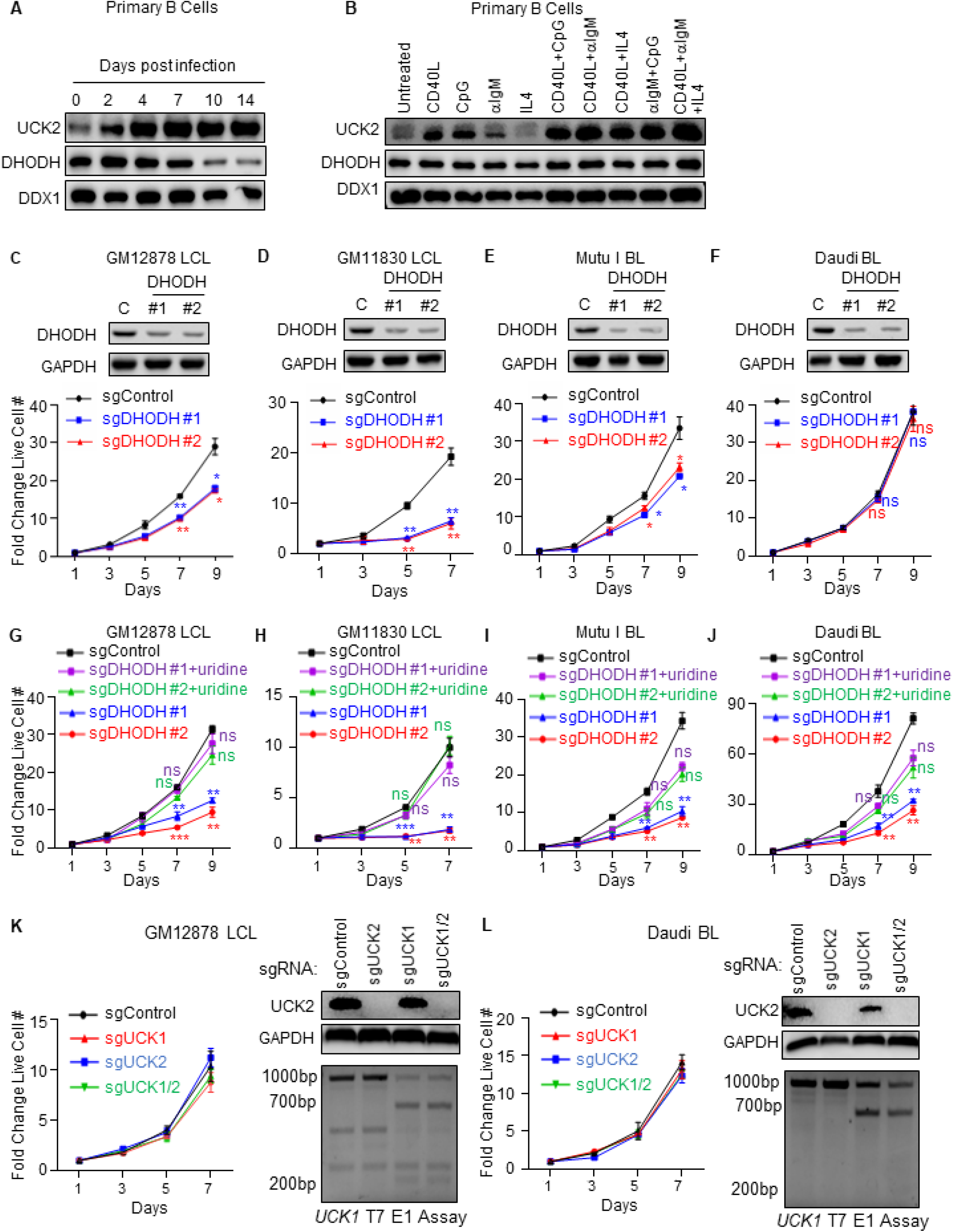
De novo pyrimidine synthesis enzyme DHODH and uridine salvage roles in EBV-transformed B-cell growth. **(A)** Immunoblot analysis of UCK2, DHODH and load-control DDX1 levels in whole cell lysate (WCL) from human CD19+ peripheral blood B-cells infected by the EBV B95.8 strain for the indicated days. (**B**) Immunoblot analysis of UCK1, DHODH and DDX1 abundances in primary human CD19+ B-cells stimulated for 24 hours by CD40L (50ng/mL), CpG (1μ M), □IgM (1μg/mL), IL-4 (20ng/mL), or combinations thereof, as indicated. **(C-F)** Immunoblot analysis (top) and growth curve analysis (bottom) of GM12878 (C), GM11830 (D), Mutu I (E) and Daudi (F) expressing non-targeting control or independent sgRNAs against *DHODH* in media containing 10% undialyzed fetal bovine serum (FBS). WCL were obtained at day 7 post sgRNA expression. **(G-J)** Growth curve analysis of GM12878 (G), GM11830 (H), Mutu I (I) and Daudi (J) expressing control or independent sgRNAs against DHODH. Cells were grown in media containing dialyzed FBS and supplemented with 200 □M uridine where indicated. **(K)** Growth curve analysis (left) and immunoblot of WCL or T7 endonuclease I (T7E1) assays of CRISPR editing of the *UCK1* locus (right) from GM12878 expressing the indicated sgRNAs. **(L)** Growth curve analysis (left) and immunoblot of WCL or T7E1 assays of CRISPR editing of the *UCK1* locus (right) from Daudi expressing the indicated sgRNAs. Blots in A-F, K and L are representative of n = 3 experiments. SEM + Mean values are shown in C-L from n=3 replicates. *, *p* < 0.05; **, *p* < 0.01; ***, *p*<0.001; ns, non-significant using one way Anova with multiple comparisons.

DHODH inhibition prevents EBV lymphomas in a humanized mouse model (24). We next used CRISPR to test the effects of DHODH depletion in LCLs or Burkitt cells. In media containing undialyzed fetal bovine serum (FBS), which contains uridine (**Figure 1a**), independent DHODH sgRNAs impaired proliferation of both LCLs and of Mutu I, but not Daudi Burkitt cells (**Figure 5c-f**). To more closely examine the ability of EBV-transformed B-cells to use uridine as a substrate for *de novo* CTP synthesis, control and DHODH edited LCL and BL were grown with dialyzed FBS lacking uridine. In this context, DHODH depletion more strongly inhibited proliferation, which was rescued by uridine add-back (**Figure 5g-j, Figure S5a-d)**. Thus, LCLs can use amino acid *de novo* or uridine salvage for CTP synthesis (**Figure 1a**).

We next used CRISPR to test the effects of UCK1 and/or UCK2 depletion on EBV-transformed B-cell proliferation. Successful CRISPR editing of UCK2 was confirmed by immunoblot, whereas UCK1 editing was confirmed by T7 E1 assay, as we were unable to find a suitable immunoblot antibody. Unexpectedly, depletion of UCK1, UCK2 or UCK1/2 did not significantly impair GM12878 LCL or Burkitt cell proliferation (**Figure 5k-l, Figure S5e**). These results suggest that LCLs and Burkitt cells can synthesize sufficient CTP *de novo* when sufficient glutamine and aspartate are available, but that they are nonetheless able to switch to uridine salvage when necessary.

### CTPS1 roles in EBV lytic DNA replication

Knowledge remains incomplete of how EBV remodels CTP synthesis pathways in support of the B-cell lytic cycle, where the viral immediate early transcription factors BZLF1 and BRLF1 induce early genes responsible for viral DNA replication and subsequently expression of viral late genes. Lytic replication results in synthesis of hundreds to thousands of EBV ∼170 kilobase genomes by the viral DNA polymerase (39). To test *de novo* cytidine roles in EBV lytic replication, control or *CTPS1* targeting sgRNAs were expressed in EBV+ Akata Burkitt cells, a model for EBV lytic replication. Following CRISPR editing, lytic replication was induced by B-cell receptor cross-linking for 24 hours. Interestingly, while CTPS1 depletion did not appreciably affect expression of the immediate early gene BZLF1 or early gene BMRF1, production of lytic EBV genomes was significantly impaired by independent CTPS1 sgRNAs (**Figure 6a**). Lytic EBV genome replication was rescued by cytidine add-back, suggesting that either the *de novo* or salvage CTP pathways can support the viral lytic cycle (**Figure 6b**). Similar results were obtained in P3HR-1 Burkitt cells, in which lytic replication was induced by a conditional *BZLF1* allele together with sodium butyrate (**Figure 6c-d**).

**Figure 6.**
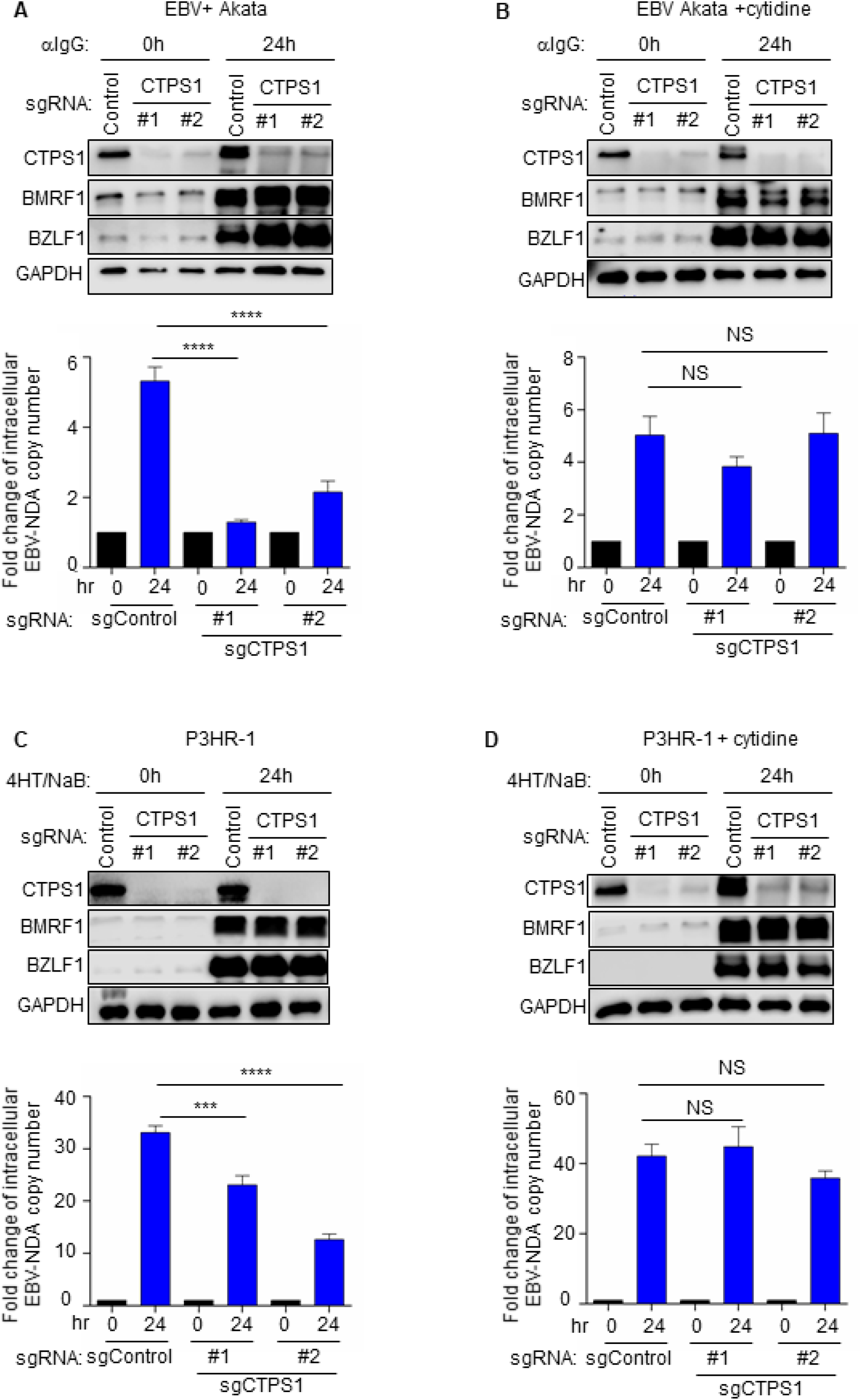
Key CTPS1 role in EBV lytic DNA replication. **(A)** Immunoblot analysis of WCE from EBV+ Akata BL expressing the indicated sgRNAs for 96 hours and mock-induced or inducef for lytic replication by □IgG (10 □g/ml) cross-linking for 24 hours. Blots for immediate early BZLF1 and early BMRF1 lytic antigens, CTPS1 and GADPH are shown. Shown below are EBV genome copy numbers determined by qPCR analysis. Values for mock-induced cells with sgControl were set to 1. Relative intracellular EBV DNA copy numbers were determined by qPCR using primers to the *BALF5* gene and were normalized for input cell number using primers to *GAPDH*. (**B**) Immunoblots and qPCR analysis of EBV genome copy number as in panel A, from cells grown in the presence of 200 □M cytidine rescue. **(C)** Immunoblot analysis of WCE from Cas9+ P3HR-1 Burkitt cells expressing a conditional 4-HT regulated BZLF1 allele and the indicated sgRNAs, mock-induced or induced for EBV lytic reactivation by addition of 400nM 4HT and 500 sodium butyrate for 24 hours. sgRNAs were expressed for 6 days. Blots for EBV-encoded wildtype BZLF1 and BMRF1 as well as for CTPS1 and GADPH are shown. Shown below are qPCR analysis of EBV genome copy number, as in A. (**D**) Immunoblots and qPCR analysis of EBV genome copy number as in panel C from cells grown in the presence of 200 □M cytidine rescue. Blots in A-D are representative of n=3 replicates. Mean + SEM values from n=3 of biologically independent replicates are shown in A-D. ****, *p*<0.0001; \*\*\**p*<0.001; NS, non-significant using one way Anova with multiple comparisons.

## Discussion

The ability of B and T lymphocytes to rapidly switch from a quiescent state into rapidly growing blasts upon antigen receptor and co-receptor stimulation underlies adaptive immunity. CTPS1 is highly induced by T-cell stimulation through the ERK pathway and is necessary for rapid T-cell proliferation and the control of acute EBV infection (23). Yet, with hypomorphic CTPS1 activity, patients often exhibit severe infectious mononucleosis, chronic active EBV and EBV+ CNS B-cell lymphomas. Since EBV relies on host metabolism pathways to convert newly infected cells into hyperproliferating blasts and to support lytic replication, these clinical observations raise the question of how individuals with CTPS1 deficiency exhibit elevated EBV viral loads and frequent B-cell lymphomas, despite hypoactivity of a key CTP generation enzyme, yet high CTP demands of B-cell growth (16, 23). Further adding to this question, classic studies indicate that CTP synthase activity is frequently upregulated in cancer (40).

We therefore tested the effects of CTPS1, CTPS2 and CTP salvage pathways to support EBV-infected B-cell growth, survival and EBV lytic replication. EBV-mediated B-cell growth transformation proceeds through a genetically encoded process that proceeds through Burkitt-like hyperproliferation and lymphoblastoid phases, whose physiology resembles the Burkitt and LCL cell lines used in this study (28, 40). LCLs are an established tissue culture model for immunoblastic lymphomas of immunosuppressed hosts. The results presented here suggest that while CTPS1 activity is important for EBV-transformed B-cell proliferation, cells can also utilize CTPS2 activity and cytidine salvage metabolism to support CTP synthesis for DNA, RNA phosphatidylcholine and phosphatidylserine needs. While knowledge of cytidine concentrations in tonsil and lymphoid tissues remains limited, robust deoxycytidine kinase activity is evident in human tonsil tissue (41).

We speculate that in patients with CTPS1 deficiency, CTPS2 compensation is insufficient for cell-mediated control of EBV infection, but is nonetheless able to support EBV+ B-cell growth, together with hypomorphic CTPS1. Given our findings that CTPS2 supports proliferation of EBV-transformed CTPS1 deficient B-cells, we speculate that CTPS2 plays important roles in the CNS lymphomas observed clinically. Consequently, CTPS1/2 may serve as a therapeutic synthetic lethal target in this setting, for instance if cytidine replacement fails to restore anti-tumor immunity. While our quantitative proteomic analyses identified that CTPS1 is approximately four times more abundant than CTPS2 in primary human B-cells undergoing EBV-mediated growth transformation (28), EBV-infected lymphomas may compensate for CTPS1 deficiency by increasing CTPS2 levels and/or activity. It would be of interest to test this hypothesis through RNAseq and enzymatic profiling of CNS lymphoma samples.

CTPS1 and CTPS2 are each highly regulated at the post-translational level, including by assembly into filaments called cytoophidium that regulate catalytic activity (42-44). Whether cytoophidium form in EBV-infected B-cells generally or with CTPS1 deficiency is unknown. Furthermore, cryo-EM studies recently identified a novel filament-based mechanism of CTPS2 regulation that stabilizes nucleotide levels (45). Filaments containing CTPS1/2 and IMPDH, the rate limiting enzyme in *de novo* GTP synthesis have also been described (46), and may play important roles in EBV-infected B-cell metabolism.

CTP salvage pathways may also play important roles in support of pathogenic B-cell roles in CTPS1 deficient patients. Cytidine is imported into the CNS by nucleoside transporters located at the blood-brain barrier (47). That CTPS1/2 deficient LCL growth could be restored by cytidine supplementation highlights the potentially important role of the CTP salvage pathway in B-cells with the viral latency III program observed in CNS lymphoma. Interestingly, since oral intake may alter CNS pyrimidine levels (47), dietary cytidine and uridine restriction could potentially have antineoplastic effects in CTPS1 deficient patients with CNS lymphoma.

Patients with CTPS1 deficiency often have elevated EBV genome copy number, which may result from latent or lytic states with impaired T-cell surveillance (48). Viral lytic gene expression has been associated with EBV lymphomagenesis (49-51). Interestingly, the DHODH chemical inhibitor teriflunomide blocks EBV lytic gene expression and DNA replication (24). Here, we found that CTPS1 knockout instead reduces EBV lytic DNA replication, without impairing immediate early and early gene expression. Consistent with a potential role in supplying CTP for lytic genome synthesis, cytidine supplementation rescued EBV lytic DNA replication, newly identifying their ability to utilize cytidine salvage metabolism. It will be of interest to define relative CTPS1 and CTPS2 roles in epithelial cell EBV lytic replication.

CTP levels increase in EBV-transforming B-cells, though to lower magnitude than other deoxynucleotide triphosphates (52). Feedback and allosteric regulation renders CTPS1 activity sensitive to the levels of all four nucleotides (22, 53). EBNA2 and MYC are important regulators of EBV-driven one-carbon metabolism and purine nucleotide synthesis in newly infected cells (28). Thus, their roles in CTPS1 induction likely coordinates pools of all four deoxynucleotides to avoid imbalances and DNA damage. Indeed, CRISPR knockout of CTPSI resulted in □□H2AX phosphorylation. While NF-□B is not required for initial EBV-driven B-cell growth (54), it is essential for LCL growth and survival, where we found a non-canonical NF-□B role in CTPS1 expression. Our findings also suggest that hyper-active MYC activity is also sufficient to drive CTPS1 expression and likely supports CTPS1 expression in Burkitt cells. By contrast, ERK and SRC have obligatory roles in robust CTPS1 upregulation upon T-cell activation that are critical for T-cell expansion (**Figure 7**). Notably, MYC has key roles in *de novo* pyrimidine nucleotide synthesis (55).

**Figure 7.**
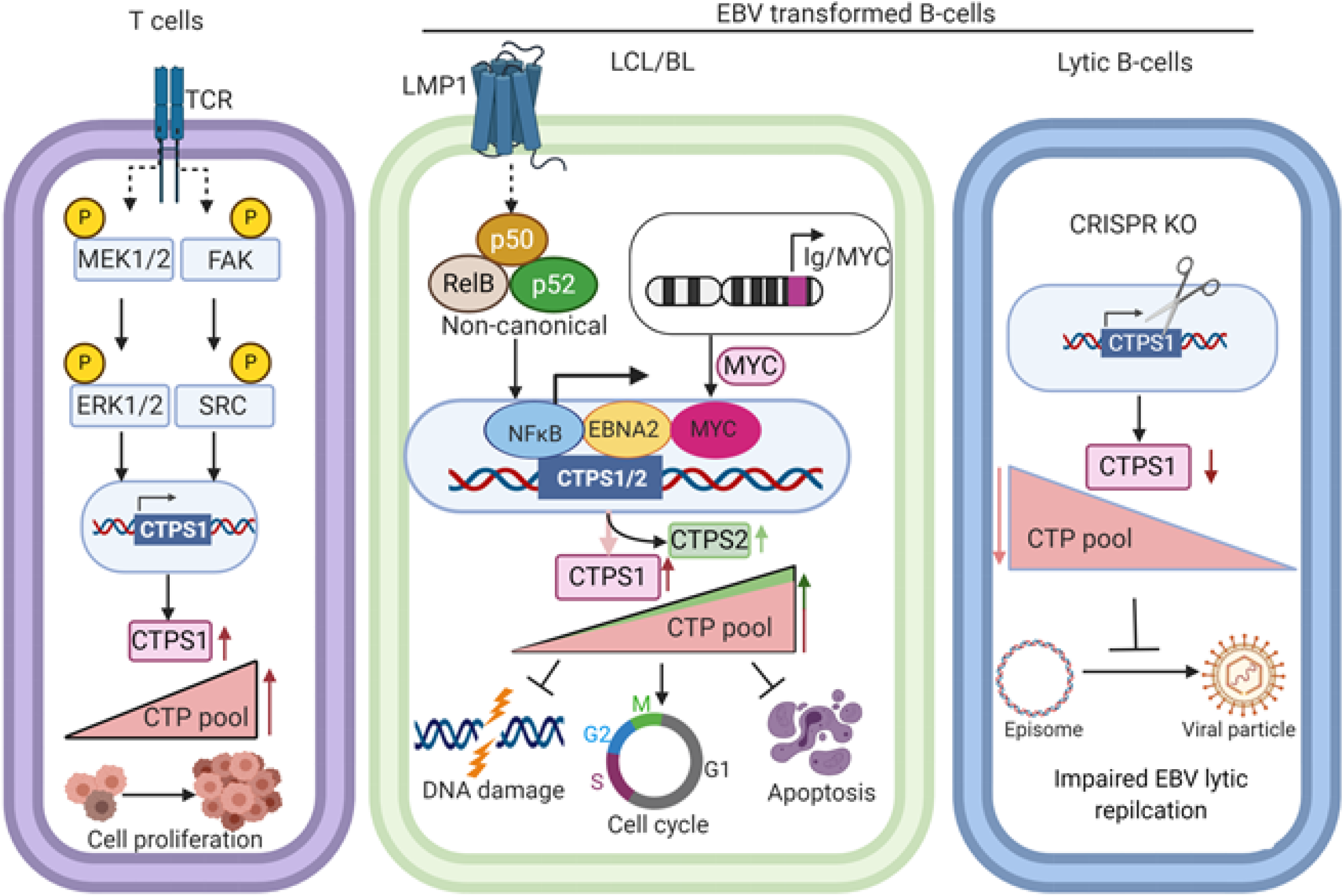
Schematic model of CTPS1 roles in T vs EBV-infected B-cells. T-cell receptor stimulation induces CTPS1 expression, at least in part through the ERK and SRC pathways. By contrast, EBV induces CTPS1 in LCLs through EBNA2, MYC and LMP1-activated non-canonical NF-□Bpathway, whereas hyper-expressed MYC has key roles in Burkitt CTPS1 induction. CTPS1 and CTPS2 activity together supply CTP pools for EBV-transformed B-cells to prevent nucleotide imbalance, DNA damage, cell cycle arrest and apoptosis. CTPS1 depletion also impairs EBV lytic DNA replication.

In summary, these studies highlight that multiple CTP synthesis pathways are operative in EBV-infected B-cells, and that each can play important roles in EBV transformed B-cell growth and lytic DNA replication. Collectively, these results highlight CTP synthesis pathways as important therapeutic targets in EBV-driven lymphoproliferative disorders.

## Methods

### CRISPR/Cas9 gene editing

CRISPR editing was performed as previously described (56). Briefly, Broad Institute pXPR-011 pXPR-515 control sgRNA, Avana or Brunello library sgRNAs, as listed in **Supplementary Table 1**, were cloned into lentiGuide-Puro (Addgene Cat#52963) or pLenti SpBsmBI sgRNA Hygro (Addgene Cat# 62205). sgRNA expressing lentiviruses were produced by 293T transfection and used to transduce LCLs or Burkitt cells with stably expressed Cas9.

### Growth Curve Analysis

Cells were normalized to the same starting concentration. Cell numbers were then quantified, using the Cell Titer Glo assays at the indicated times.

### Recombinant EBV

EBV B95-8 virus was produced from B95-8 cells with conditional BZLF1 expression. The P3HR-1 EBV strain was produced from P3HR-1 cells with conditional BZLF1 and BRFL1 expression. UV irradiation of B95-8 virus supernatants was performed at a cumulative intensity of 3J per square centimeter on ice, to prevent heat-induced virus degradation. The quantitation of equal infection of B95-8, UV B 95-8 and P3HR1 was performed as previously described (57).

### Statistical Analysis

Data are presented as means ± standard errors of the means. Data were analyzed using analysis of variance (ANOVA) with Sidak’s multiple-comparison test or two-tailed paired Student’s *t* test with Prism7 software. For all statistical tests, a *P* cutoff value of <0.05 was used to indicate significance.

## Acknowledgements

This work was supported by NIH RO1s AI137337 and CA228700 (BEG) and a Burroughs Wellcome Career Award in Medical Sciences and CA047006 to BZ. We thank Jaap Middeldorp for generously sharing multiple anti-EBV antibodies.

## Compliance with ethical standards

Conflict of interest BG receives research funding from Abbvie unrelated to this study.

## Author contributions

J.L. performed the experiments. C.W, R.G., S.Y., B.Z. provided technological assistance. R.G. performed bioinformatic analysis. J.L, R.G. and B.E.G. prepared the manuscript. B.E.G supervised the study.

## Supplementary Figure Legends

**Figure S1. EBV-induced *de novo* pyrimidine synthesis pathway expression**.

**(A-F)** CAD (A), DHODH (B), UMPS (C) CMPK1 and CMPK2 (D), CTPS1 and CTPS2 (E), NME1 and NME2 encoded NDPK (F) relative protein abundances (top) and mRNA abundances (bottom) over the first 28 days of peripheral blood CD19+ B-cell infection by EBV B95.8 at a multiplicity of infection of 0.1. Mean + SEM values from n=4 tandem-mass-tag-based proteomic and n=3 RNAseq replicates were used for these analyses and were obtained from recently published datasets (28, 29). *, *p*<0.05; **, *p*<0.01; ***, *p*<0.001; ****, *p*<0.0001; ns, non-significant, using one-way Anova with multiple comparisons.

**Figure S2. CTPS1 roles in EBV-transformed B-cell growth and survival**.

(**A-B**) Immunoblot analysis (top) and growth curve analysis (bottom) of GM11830 LCLs (A) and Mutu I BLs (B) expressing control or independent sgRNAs against CTPS1. WCL were obtained at day 7 post sgRNA expression.

**(C)** FACS analysis of 7-AAD uptake in GM11830 LCL (left) or Mutu I BL (right) expressing control or CTPS1 sgRNAs for 7 days.

(**D**) Median + SEM values from n=3 replicates, as in C.

**(E)** Propidium iodide cell cycle analysis of GM12878 LCL, GM11830, Daudi BL or Mutu I BL expressing control or CTPS1 sgRNAs at Day 9 post sgRNA expression.

(**F**) Median + SEM values from n=3 replicates, as in E. Percentages of cells in G2/M, S, G0/G1 or with <2n DNA content indicative of cell death are shown.

**(G)** Immunoblot analysis of WCL from GM11830 (left) or Mutu I cells (right) expressing control or CTPS1 sgRNAs for 7 days.

Blots in A and G are representative of n = 3 experiments. SEM + Mean values are shown in A, D and F from n=3 replicates. *, *p* < 0.05; **, *p* < 0.01; ***, *p*<0.001; ****, *p*<0.0001; ns, non-significant using one-way Anova with multiple comparisons.

**Figure S3. Analysis of CTPS2 roles in EBV-transformed B-cell growth**.

**(A)** DepMap CRISPR screen dependency scores (31) for sgRNAs targeting CTPS1 (left) and CTPS2 (right) across cell lines from the indicated cancer cells of origin. Each oval represents the DepMap screen value for a cell line from the tissue indicated at right. Negative values indicates selection against the sgRNAs target the gene over a 21 day growth and survival screen. Values <-1 (red vertical line) indicates cell line dependency on the targeted gene for growth or survival.

(**B-C**) Immunoblot analysis (top) and growth curve analysis (bottom) of GM11830 LCL (B) or Mutu I BL (C) expressing control or independent CTPS2 sgRNAs for 7 days.

**(D)** FACS analysis of 7-AAD uptake in GM12878 (left) or Daudi cells (right) expressing control or CTPS2 sgRNAs for 7 days.

**(E)** Median + SEM values from n=3 replicates, as in D, of 7-AAD uptake FACS analysis in GM12878, GM11830, Daudi or Mutu I cells.

**(F)** Mean + SEM values of FACS propidium iodide cell cycle analysis of GM12878 (left) or Daudi (right) expressing control, CTPS2, CTPS1 or CTPS1/2 sgRNAs for 7 days. Significantly differences from sgControl values are indicated.

Blots in B and C are representative of n = 3 experiments. SEM + Mean values are shown in B-F are from n=3 replicates. *, *p* < 0.05; **, *p* < 0.01; ***, *p*<0.001; ****, *p*<0.0001; ns, non-significant using one-way Anova with multiple comparisons.

**Figure S4. Profiling of EBV and immune receptor stimulation effects on UCK1, UCK2 and DHODH expression in primary B-cells**.

**(A)** qPCR analysis of UCK1 and UCK2 mRNA levels in human peripheral blood CD19+

B-cells at the indicated day post infection by B95.8 EBV at MOI=0.1. Day 0 values were set to 1.

(**B**)UCK1, UCK2 and DHODH relative protein abundances from published analyses of tandem-mass-tag-based proteomic analysis at rest and at nine time points after EBV B95.8 infection of human peripheral blood CD19+ B-cells at a multiplicity of infection of 0.1 (28).

**(C)** qPCR analysis UCK1, UCK2 or DHODH mRNA levels in human peripheral blood CD19+ B-cells stimulated for 24 hours by Mega-CD40L (50ng/mL), CpG (1μM), □IgM crosslinking (1μg/mL), IL-4 (20ng/mL) or combinations thereof, as indicated.

SEM + Mean values are shown in A-C from n=3 replicates. *, *p* < 0.05; **, *p* < 0.01; ***, *p*<0.001; ****, *p*<0.0001; ns, non-significant using one-way Anova with multiple comparisons.

**Figure S5. Analysis of DHODH, UCK1/2 and uridine salvage metabolism roles in EBV-transformed B-cell growth**.

**(A-B)** Growth curve analysis of GM11830 (A) or Mutu I (B) expressing control or DHODH sgRNAs and puromycin selected for 48 hours, then grown in media containing uridine-depleted 10% dialyzed fetal bovine serum, in the absence or presence of 200 □M uridine supplementation. Immunoblot analysis of WCL from cells with the indicated sgRNAs are shown to the right.

**(C)** FACS analysis of 7-AAD abundances in GM12878, GM11830, Mutu I or Daudi expressing the indicated sgRNAs for 7 days and grown in media containing uridine-depleted, dialyzed FBS in the absence or presence of 200□M uridine supplementation.

**(D)** Mean + SEM values of FACS analysis of n=3 replicates, as in C.

**(E-F)** Growth curve analysis (left) and immunoblot of WCL for UCK2 and GAPDH load control or *UCK1* locus T7E1 assay results (right) of GM11830 (E) or Mutu cells (F) expressing the indicated sgRNAs.

Blots in A, B, E and F are representative of n = 3 experiments. SEM + Mean values are shown in A, B, D, E and F from n=3 replicates. *, *p* < 0.05; **, *p* < 0.01; ***, *p*<0.001; ns, non-significant using one-way Anova with multiple comparisons.

